# Clearing-induced tisssue shrinkage: Novel observation of a thickness size effect

**DOI:** 10.1101/2020.04.20.051771

**Authors:** R.C.M. Vulders, R.C. van Hoogenhuizen, E. van der Giessen, P.J. van der Zaag

**Affiliations:** Philips Research Laboratories, High Tech Campus 11, 5656 AE Eindhoven, The Netherlands; Fontys University of Applied Sciences, Rachelsmolen 1, 5612 MA Eindhoven, The Netherlands; Zernike institute for Advanced Materials, University of Groningen, Groningen, The Netherlands

## Abstract

The use of clearing agents has provided new insights in various fields of medical research (developmental biology, neurology) by enabling examination of tissue architecture in 3D. One of the challenges is that clearing agents induce tissue shrinkage and the shrinkage rates reported in the literature are incoherent. Here, we report that the shrinkage for a widely-used clearing agent decreases significantly with increasing sample size, and report an analytical description.

---

Solvent-based tissue clearing is a widely used methodology to render an otherwise opaque sample optically transparent. Using tissue clearing in combination with various optical imaging techniques enables the study of various organs and organism, such as mouse brains^1–3^, larvae and spinal cords^4^ as well as tumours including their (micro)environment^5^ such that the overall structure and interaction with the neighboring tissue structures can be studied and understood. This has been important for the advancing understanding in neurology and oncology.

Typically in tissue clearing, samples are immersed in various solvents and incubated to achieve complete permeation of the solvent and clearing of the specimen, as demonstrated already over a century ago^6^. Tissue clearing is achieved by consecutive steps of: fixation, dehydration, de-lipidation (using solvents such as paraformaldehyde, methanol) and finally refractive index matching using a clearing agent (such as mixture of benzyl-alcohol and benzyl-benzoate (BABB)) ^4,7^. Solvent-based clearing strategies all share the disadvantage that during the different processing steps alterations in the overall sample size and possibly architecture may occur, due to shrinkage of the tissue^8^. Alternatives to solvent-based clearing procedures are simple immersion in an aqueous solution with sugars or using hyperhydration^9^. Yet, in view of the good clearing they produce, solvent-based clearing methods based on BABB^1,4,10^ or derived agents^11^ play a prominent role in the field.

Strikingly, upon detailed examination of the literature about the extent of tissue shrinkage induced by clearing agents, a very confusing picture emerges, with various sources pointing to previous literature, which again refers to others. In the end, most sources point to the work by Ertürk *et al*. who reported that the tissue shrinks isotropically in all dimensions by 21%, resulting in a 51 % reduction of the 3D volume^4^. However, one of the more recent paper reports different amounts of shrinkage for different tissues (e.g. 55 % for mouse brains, 30 % for hippocampus and cortex layers, and 10.7 % for a mouse torso)^11^. Given that a number of groups are extending the study of tissue analysis in 3D to that of human biopsies, using various means of optical microscopy yet all relying on tissue clearing^10,12,13^, it becomes important to know the extent of tissue shrinkage caused by the use of clearing agents.

In this study, we systematically determined the amount of shrinkage for different tissue types in each of the consecutive steps needed to clear tissue, by measuring the diameter of cylindrical biopts. We have investigated this for BABB, being a frequently used and prominent example of a clearing agent. Sampling was performed using sampling needles or dermal biopsy punchers with different inner diameters in order to obtain samples with various initial diameters. All samples were processed according to the same protocol. The samples were washed with phosphate buffered saline (PBS) and subsequently fixed overnight in a 4 % paraformaldehyde solution. The next day the samples were dehydraded using a graded methanol washing series (in steps of 25 %) for 1 hour each. The last two processing steps were a 50 % methanol/50 % BABB step, after which the final clearing in 100 % BABB was performed (see **Fig. 1a** and for more details the online methods section). Microscopy images were acquired for each sample. **Fig. 1** shows an example for a rat liver tissue. The consecutive panels display the same tissue sample after each step in the dehydration- and clearing procedure. The shrinkage of tissue is well visualized across the different solvents used in the procedure. The diameter was recorded and compared to the initial tissue diameter D (see for more detail the online methods section and **Supplementary Fig. 2).** The reduction in diameter, ΔD, for all used tissues and different initial diameters are given in **Fig. 2**. In **Fig. 2a** the diameter change is plotted versus initial diameter for three different tissues; two of animal origin (rat liver and rat spleen) and one of human tissue origin (prostate). The reduction in diameter seems to level off with increasing initial tissue diameter. Note that after the procedure all samples were completely cleared, see **Fig. 1** (i). To verify the levelling-off of the shrinkage for the thickest samples, the experiment for the thickest rat liver sample was repeated with the time of each individual processing steps extended from 1 to 3h. At 407 ± 34 µm this did not yield a different result in the observed tissue shrinkage.

**Fig. 1.**
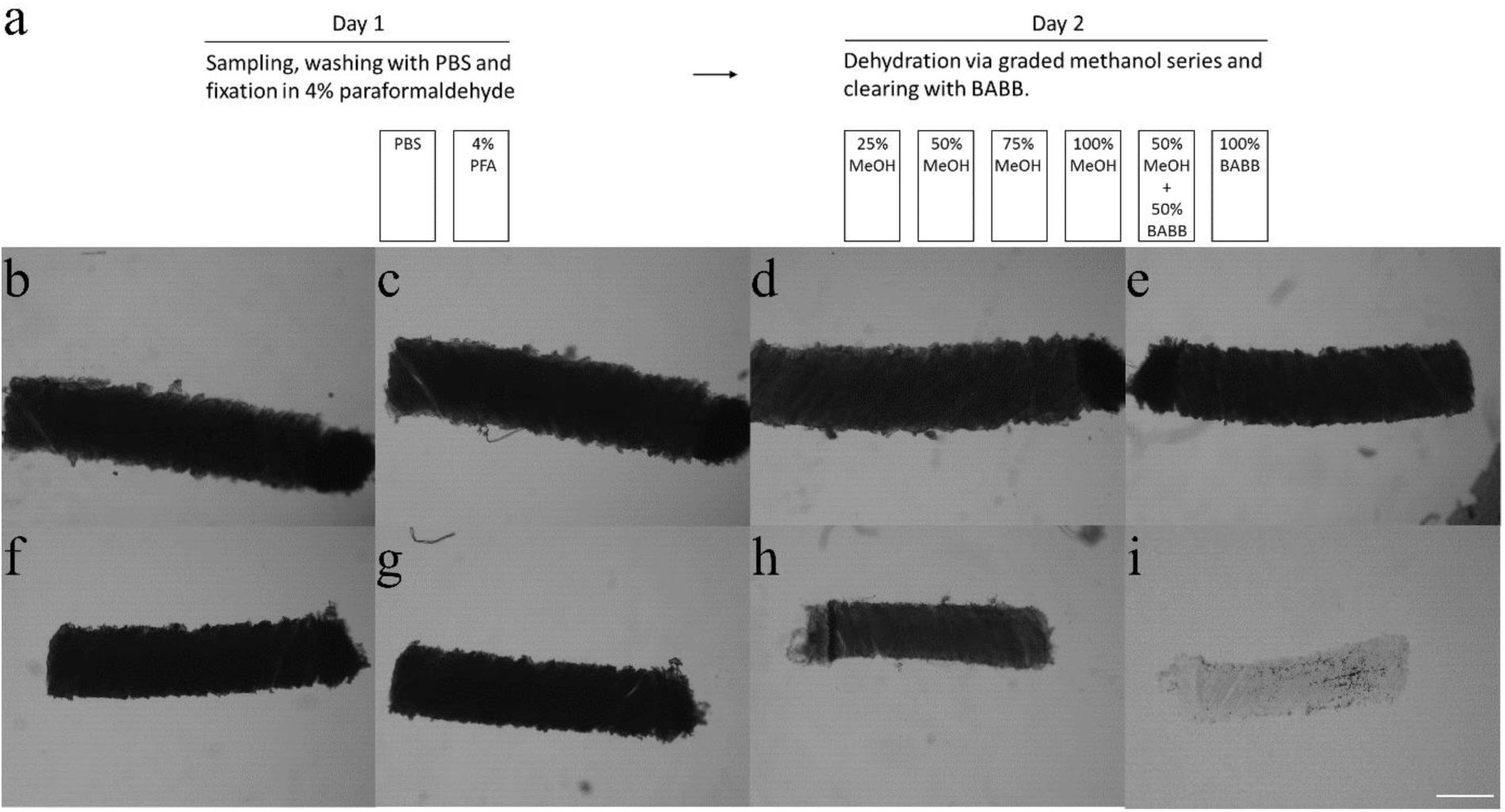
Optical microscopy images acquired after each consecutive process step of the clearing protocol for a rat liver sample. Panel **(a)** gives a schematic representation of the procedure. Panel **(b)** shows the sample immediately after sampling, in **(c)** after overnight fixation, **(d-g)** after the graded methanol series, **(h)** after the 50% methanol / 50% BABB step and **(i)** after the final clearing step using the clearing solvent BABB. All images were acquired with the same settings. For visualization purposes the magnification was kept at 4x in all images. The scale bar depicted in panel (**i**) represents 500 µm for all panels.

**Fig. 2.**
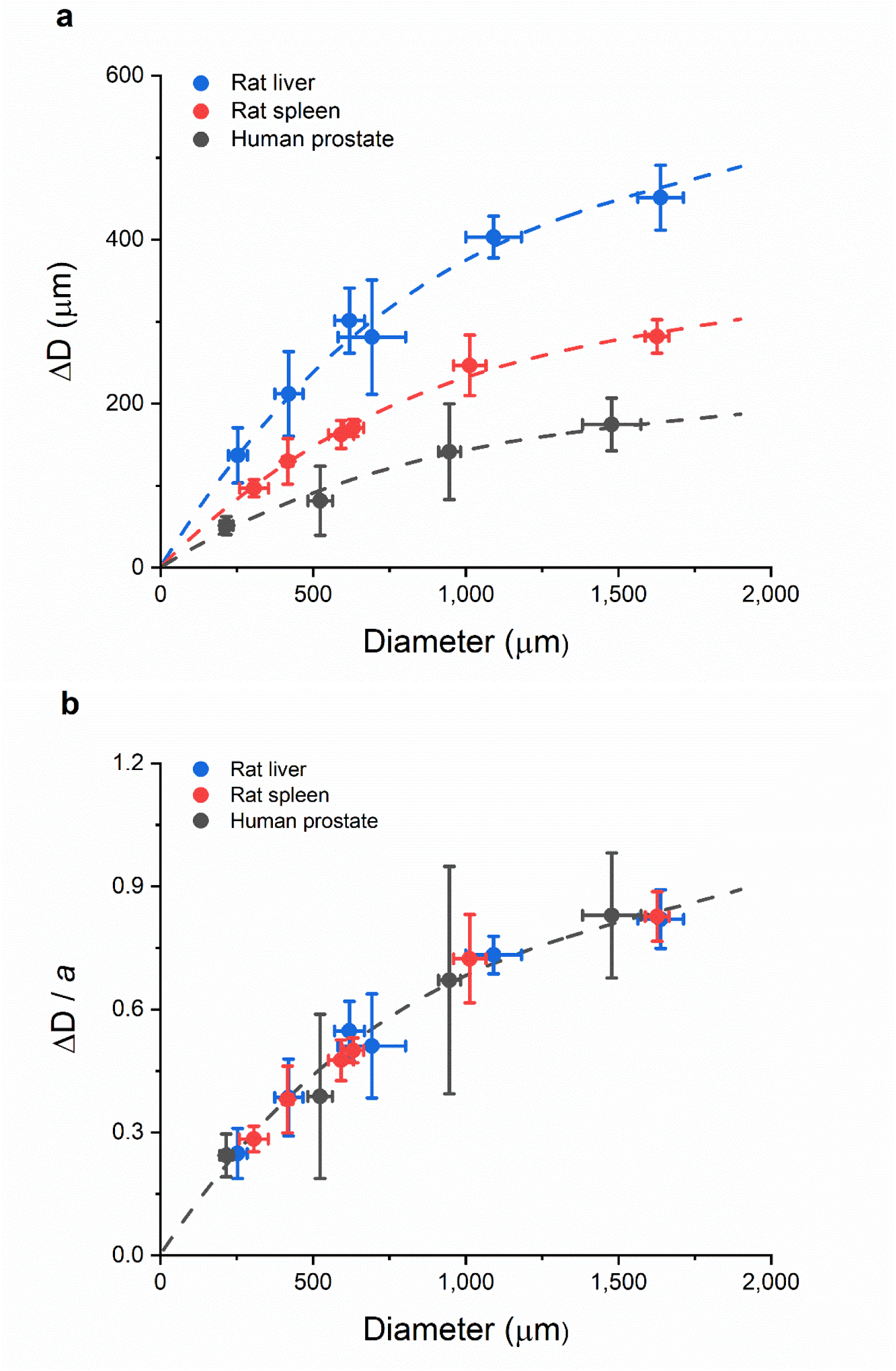
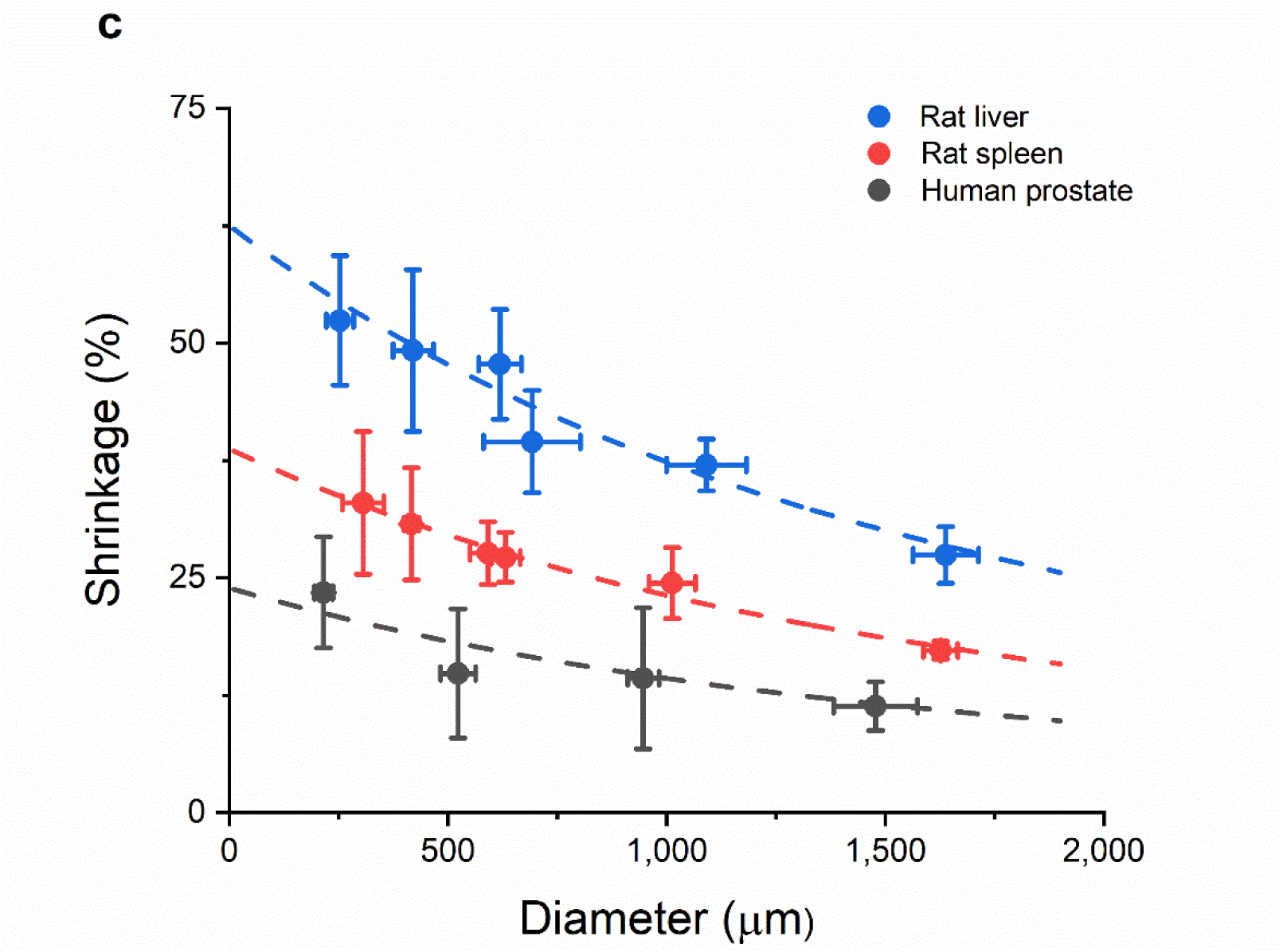
**(a)** Plot of the measured diameter change (ΔD) in µm for 3 different tissues: rat liver (blue), rat spleen (red) and human prostate (black) as a function of the initial sample thickness D due to sampling with different needle diameters. The dotted lines are a fits to the Eq. (1) see text. **(b)** Plot of the diameter change normalized by the maximum change *a* as a function of the initial sample thickness due to sampling with different needle diameters. The line in the figure is a fit based on Eq. (1). Note that the relative tissue shrinkage for all three tissue types fall on a universal curve. **(c)** The tissue shrinkage S=ΔD/D as a function of diameter for each of the 3 tissues examined. The dotted lines is based on a fit to Eq. (2). Note that the shrinkage for all three tissue types decreases with increasing D and thus is not a constant for a tissue.

The trend in the diameter change can be described with the following function:

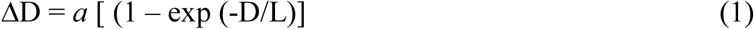

in which *a*_*i*_ is a tissue dependent factor giving the maximum shrinkage in the limit that the tissue diameter, D → ∞, and L is a length scaling factor. As shown in **Fig. 2a**, this expression fits all three curves very well for the same value L = 880 µm and for tissue dependent values *a* = 550 µm (liver), 340 µm (spleen) and 220 µm (prostate), respectively. Not unexpectedly, the magnitude of the shrinkage depends on tissue type, consistent with recent literature reports^11^. Moreover, we note that the *a*-values, which reflect how sensitive the tissue is to shrinkage, are in line with the general trend in fat percentage of these tissues^14^, which is consistent with the fact that BABB clearing includes the removal of lipids^9^. This observation immediately suggests that normalization of the diameter change by the tissue dependent factor *a*_*i*_ should give universal behavior, which is confirmed by **Fig. 2b**.

The reduction in diameter of the samples is the direct read-out of the experiment, but is not a physically meaningful characteristic. The shrinkage S = ΔD/D is meaningful and is plotted in **Fig. 2c** (in %, as is common in the literature). This figure shows clearly that the shrinkage, S, is *not* constant for a given tissue but depends on the sample diameter D in a manner which can be simply derived from Eq. (1) to be:

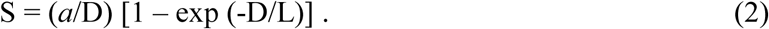

The curves in **Fig. 2c** are based on Eq (3) and the parameter values given above. A series expansion of the exponential in Eq. (2) for small values of (D/L),

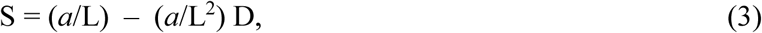

reveals that the tissue shrinkage varies linearly with D for small samples and that S = *a*/L in the limit that D → 0.

Our experimental data present, first of all, a caveat to the community that, in contrast to the existing literature, the amount of tissue shrinkage by BABB clearing depends in addition to the tissue also on the size of the sample. Secondly, this work shows that it is possible to establish a closed-form empirical description of size dependent shrinkage, which will be instrumental for a quantitatively correct interpretation of images of BABB-cleared biopsies in the clinical practice^10,12,13^. At this point, we do not know the physical mechanism responsible for the observed size dependent clearing. Yet, it may be sought in the diffusion and reaction aspects of the clearing process itself. The experimental data presented here provides a detailed reference for the validation of a full microscopic and predictive model of the clearing process.

In conclusion, we have shown that the tissue shrinkage using the commonly used BABB clearing method is not fixed but depends on the sample size. For three different tissue types this dependence can be described by the empirical relation given by Eq. (3), at least up to sample sizes of 1600 µm. Further work to examine this for other clearing agents is being undertaken.

## METHODS

### SAMPLE PROCESSING

Sampling was performed using needles with different inner diameters: 300, 550, 800, 900 and 1200 µm (BD, New Jersey, USA). Needles and Syringes (BD, New Jersey, USA) were filled with sterile phosphate buffered saline (PBS, Merck, Darmstadt, Germany) prior to puncturing, this to release the samples more conveniently. The 1.6 mm sampling was performed by using a dermal biopsy puncher (Integra, Miltex, York, Pennsylvania, USA, inner diameter 1.67 mm). Biopsies were taken from frozen samples stored at – 20 ^0^C. Upon puncturing, the samples were transferred into homemade solvent resistant containers at room temperature (**Supplementary Fig. 1)**. Briefly, a dual barrel piston container (Nordson EFD, Bedfordshire, England) was cut and fastened on a coverslip (Menzel-glaser, Thermo Scientific, Waltham, Massachusetts, USA) with a 2 component epoxy adhesive (UHU, Bühl/Baden, Germany); upon mounting the adhesive was left overnight to solidify.

Upon deposition the samples were washed 2 times 15 minutes with PBS prior to overnight fixation with 4% paraformaldehyde (PFA, Merck, Darmstadt, Germany). Washing steps and incubation took place under agitation at 4 °C unless stated otherwise. Subsequently the samples were cleared with a solvent based methodology^1^, with slight modifications. First the samples were dehydrated via a graded methanol (MeOH) series (Fisher Scientific, Waltham, Massachusetts, USA) starting at 25% and in incremental steps of 25% up to 100% MeOH. After dehydration, samples were taken up in a 1:1 (v/v) mixture of 100% MeOH and a 2:1 (v/v) mixture of benzyl alcohol (Sigma Aldrich, Saint Louis, USA) and benzyl benzoate (Sigma Aldrich, Saint Louis, USA) resulting in a 50% MeOH - 50 % BABB mixture. Finally the samples were taken up in 100% BABB. All incubation steps were carried out for 1 hour.

Samples used in this study were obtained from redundant material and consisted of rat spleen and rat liver. Human prostate samples were commercially purchased (Proteogenex, Inglewood, California, USA). For all diameters, 5 independent samples for the rat tissue were taken, while from the human prostate tissue 3 independent samples were taken. To follow the clearing procedure microscopic images were acquired after each incubation step with a microscope fitted with a camera (Leica, Wetzlar, Germany). To determine the shrinkage of the tissues during the clearing procedure, the diameters were determined using the scale bar function in Leica application suite (LAS) software. This was done at 6 different positions for each sample after each consecutive step in the clearing protocol.

### MEASUREMENT OF SHRINKAGE

The tissue diameter was measured after the sample was taken from the biopsy needle, as we found that the needle diameter is not equal to the initial sample diameter. Moreover, this difference depends on tissue type. Subsequently, after each of the 7 processing steps in the clearing protocol, the sample diameters were measured. For each needle diameter, 5 samples were taken, except for human prostate tissue in which 3 samples were taken for each needle diameter. At all stages, the diameter of each sample was measured at 6 points spread along the length of the entire sample as shown in **Supplementary Fig. 2**, using a scale bar available in the microscope (Leica application suite (LAS) software, version 090-135.001, Wetzlar, Germany). Thus a dispersion of measuring points and diameters was obtained, ensuring that the entire tissue is being measured for shrinkage and not just a particular part. In summary, for each data point in **Fig. 2**, for each sample its average diameter was determined from 6 measurements and a total of 5, or for prostate 3, samples were averaged.

## AUTHOR CONTRIBUTIONS

PJZ conceived the project, PJZ and RV designed the experiments. RV and RH performed the experiments and measurements. Data analysis and interpretation was performed by all authors. PJZ, RV, and EG prepared the manuscript. All authors read and approved the final version of the manuscript.

## SUPPLEMENTARY FIGURES

**Supplementary Figure 1.**
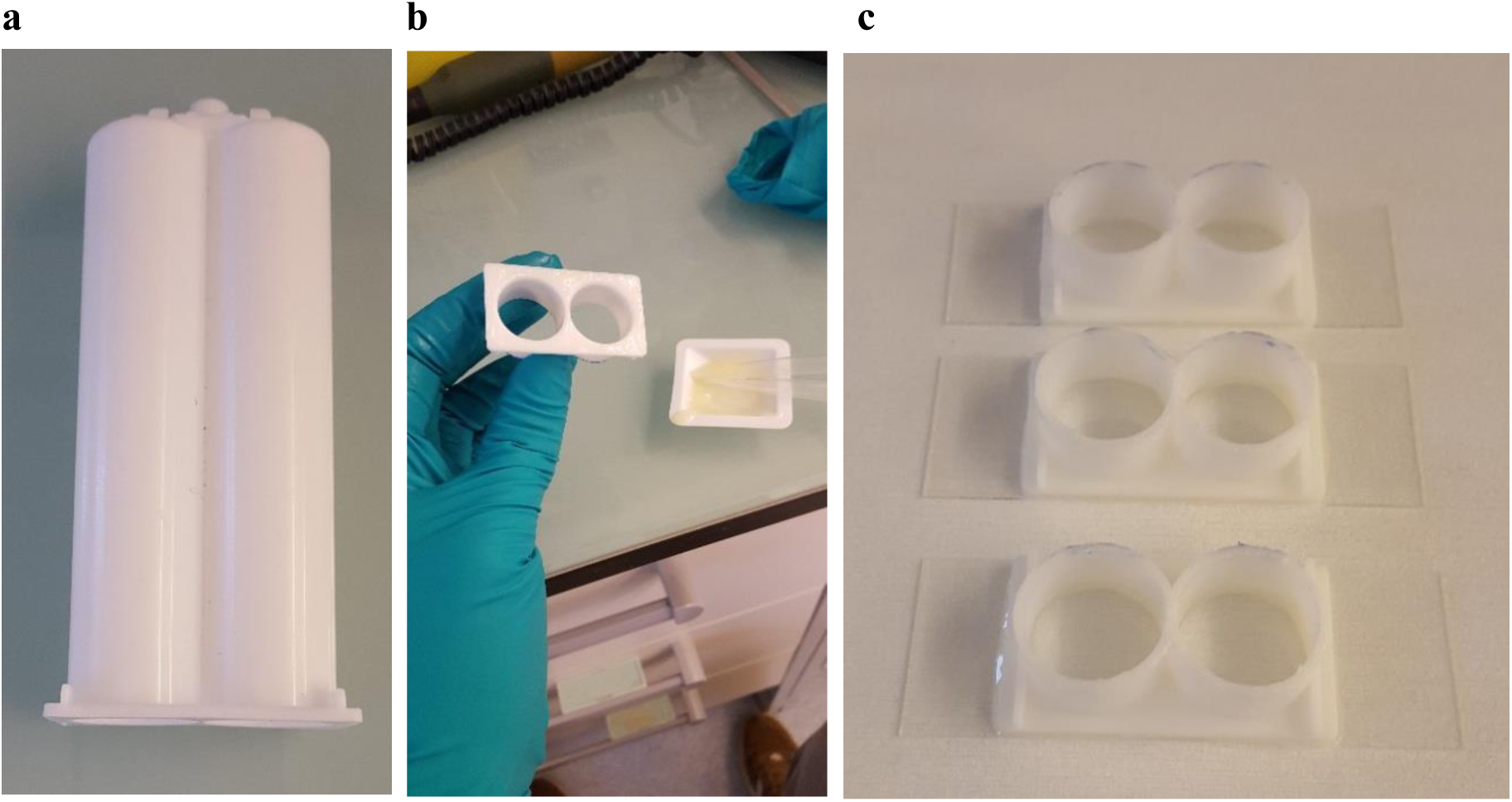
Preparation of solvent resistant containers. Depicted in **(a)** a dual barrel syringe container used to generate the solvent resistant container, **(b)** shows the assembling of the modified piston barrel onto a coverslip and **(c)** shows the completed container which can conveniently be used for imaging and solvent changes while minimizing manual handling of the samples.

**Supplementary Figure 2.**
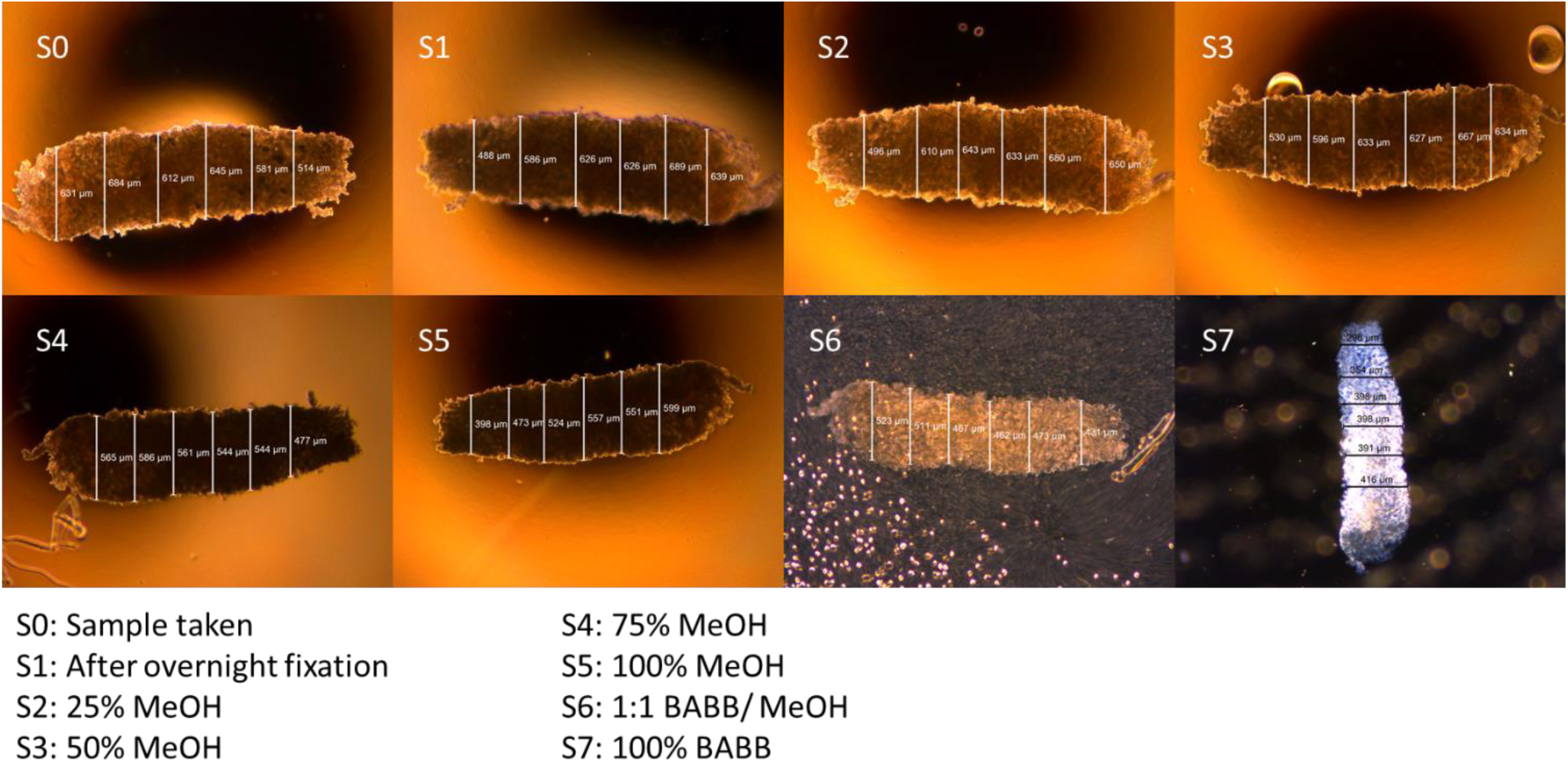
Measurement of tissue diameters during the 7 processing steps in the tissue clearing procedure. Shown is the measurement of the diameter of a single tissue sample, rat liver tissue in this particular example, for each of the processing steps (labelled Si). This figure shows the variation in diameter over the sample and the need to measure at several (in total 6) points, along the sample. The legend lists the solvents in which the tissue was incubated during the 7 process steps.

